# Probing the stability and interdomain interactions in the ABC transporter OpuA, using single-molecule optical tweezers

**DOI:** 10.1101/2023.09.01.555951

**Authors:** Lyan van der Sleen, Jan Stevens, Siewert J. Marrink, Bert Poolman, Kasia Tych

**Affiliations:** Department of Biochemistry, Groningen Biomolecular Sciences and Biotechnology Institute, University of Groningen, Nijenborgh 4, 9747 AG Groningen, The Netherlands; Molecular Dynamics Group, Groningen Biomolecular Sciences and Biotechnology Institute, University of Groningen, Nijenborgh 7, 9747 AG Groningen, The Netherlands; Chemical Biology group, Groningen Biomolecular Sciences and Biotechnology Institute, University of Groningen, Nijenborgh 7, 9747 AG Groningen, The Netherlands

**Keywords:** integral membrane proteins, ABC transporter, inter-domain interactions, optical tweezers, substrate-binding domains, single-molecule analysis

## Abstract

Transmembrane transporter proteins are essential for the maintenance of cellular homeostasis and, as such, are key drug targets. Many transmembrane transporter proteins are known to undergo large structural rearrangements during their functional cycles. Despite the large amount of detailed structural and functional data available for these systems, our understanding of their dynamics and therefore how they function is generally limited. We introduce an innovative approach which enables us to directly measure the dynamics and stabilities of inter-domain interactions of transmembrane proteins using optical tweezers. Focusing on the osmoregulatory ABC transporter OpuA from *Lactococcus lactis*, we examine the mechanical properties and potential interactions of its substrate-binding domains. Our measurements are performed in lipid nanodiscs, providing a native-mimicking environment for the transmembrane protein. The technique provides high spatial and temporal resolution and allows us to study the functionally-relevant motions and inter-domain interactions of individual transmembrane transporter proteins in real-time in a lipid bilayer.

## Introduction

Biological membranes consist of a lipid bilayer and function as a semi-permeable barrier to separate the interior and surrounding environment of a cell or organelle. Small apolar molecules such as weak acids and bases can often diffuse through this barrier with the concentration gradient. However, the majority of molecules (nutrients, ions, cofactors) require transport proteins to facilitate passage across the membrane, and in many cases these systems enable transport against the concentration gradient. Membrane transport proteins are therefore crucial to the maintenance of cellular homeostasis. Compared to soluble proteins, membrane proteins are more difficult to study as they require a lipid-mimicking environment in order to stay correctly folded and functional. Despite the challenges, the study of domain-domain interactions and (mis)folding is important in developing our understanding of how these proteins function.

Optical tweezers have shown to be a powerful tool for investigating the mechanical properties of individual proteins^1–4^. Making use of a highly focused laser beam, micron-sized particles can be captured and manipulated with a sub-nm and sub-pN accuracy. With this technique, unfolding pathways^5–9^ and inter-domain dynamics^10–12^ of soluble proteins have been studied on the single-molecule level. Distance changes induced by domain-domain interactions have been resolved as accurately as to a few nm. Additionally, studies have been performed on the reversibility of folding of relatively simple membrane proteins in a bicelle using magnetic tweezers^13,14^. However, comparable studies on more complex transmembrane proteins are absent in literature^15^.

In this study, we have examined the unfolding pattern and potential interactions of the substrate-binding domains (SBDs) of the transmembrane protein OpuA, which is part of the ATP-binding cassette (ABC) transporter superfamily^16^. OpuA and homologues are key proteins in regulating cell-volume in hypertonic conditions in bacteria and archaea. This is achieved by transporting compatible solutes, like glycine betaine, into the cell to restore turgor^17,18^.

OpuA from *Lactococcus lactis* is a tetrameric protein that is composed of two identical ATP-binding/hydrolyzing subunits (OpuAA) and two identical subunits responsible for substrate-binding and translocation (OpuABC) (**Figure 1A**). OpuAA consists of a water-soluble nucleotide-binding domain (NBD) located at the cytosolic side of the transmembrane domain (TMD)^19^. Two tandem accessory cystathionine-β-synthase (CBS) domains are fused C-terminally to the NBD, which can bind the second messenger cyclic-di-AMP^20,21^. OpuABC contains the SBD fused C-terminally to the TMD. Upon binding of the ligand glycine betaine, the SBD closes^22^. When the substrate receptor interacts with the TMD, the SBD opens again to release the substrate into TMD for transport across the membrane^21^.

**Figure 1.**
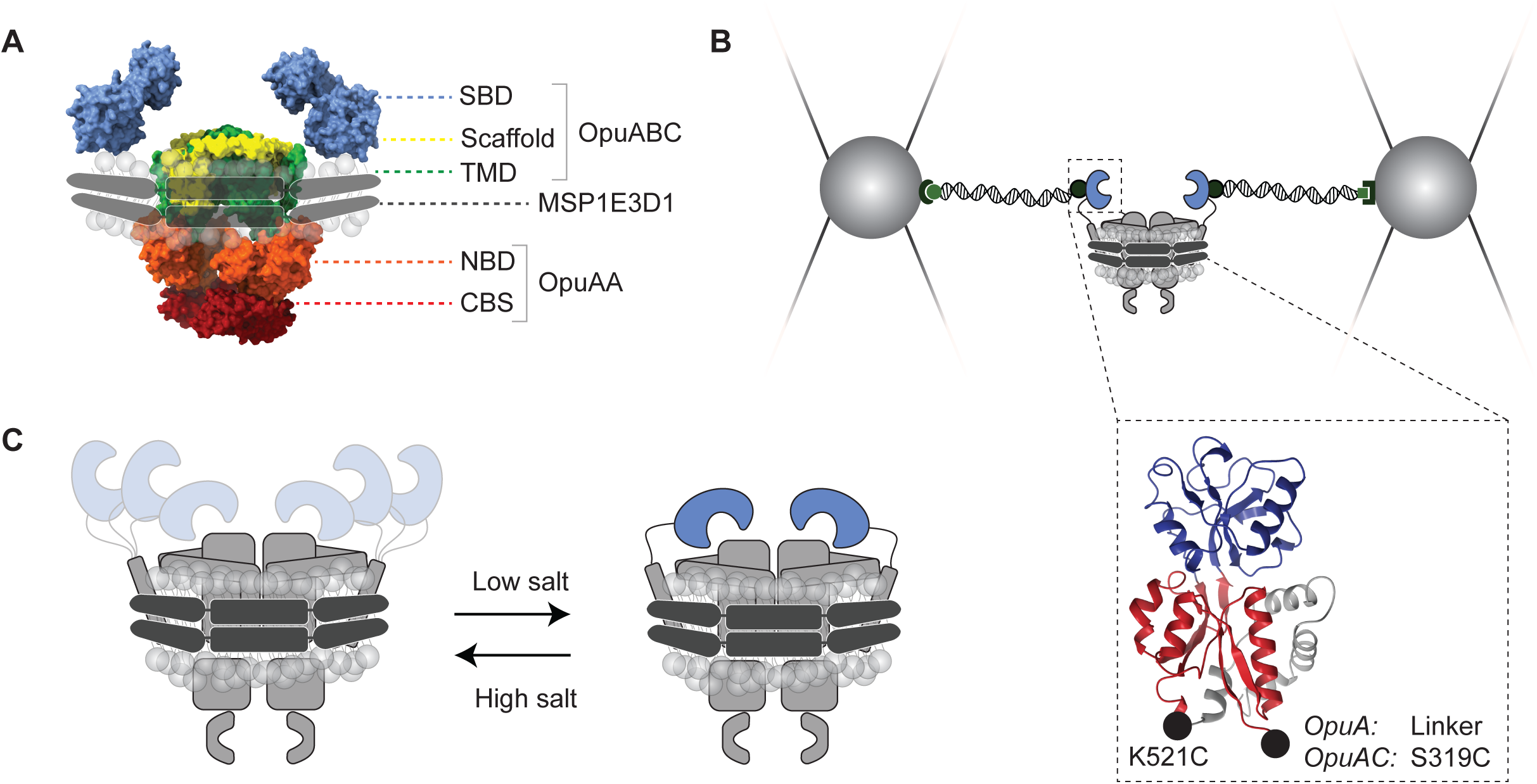
Domain organization of OpuA and experimental set-up. (**A**) Domain organization of the protomers of OpuA in lipid nanodiscs using the scaffolding protein MSP1E3D1. (**B**) Experimental set-up where silica-beads are caught by the laser and the protein is tethered between the beads with double-stranded DNA handles. Figure is not drawn to scale. Zoom-in: crystal structure (PDB: 3L6G) of the SBD and attachment points of the handles. Both OpuA and OpuAC (isolated SBD) have been studied by optical tweezers. (**C**) A proposed interaction of the SBDs. The SBDs are more flexible at high ionic strength.

Transport of the substrate glycine betaine is fully coupled and fueled by the hydrolysis of ATP^21,23,24^, which only occurs at relatively high internal ionic strength^20,24,25^. The gating of OpuA by ionic strength is mediated by interactions of the protein with anionic lipids on the cytoplasmic surface of the membrane^24,26,27^. A cationic patch (helix-turn-helix motif) in the NBD is thought to be responsible for sensing the ionic strength by locking the NBDs to the negatively charged membrane^21,26,27^. An increase in ionic strength screens the electrostatic interactions of the cationic patch with the membrane, which activates the transport. Single-molecule Förster resonance energy transfer microscopy (smFRET) and cryo-electron microscopy (cryo-EM) results have implied an interaction between the two SBDs, which is especially prominent in low ionic strength conditions^28^ (**Figure 1C**). The relative orientation of the interacting SBDs could not be deduced based on smFRET and cryo-EM results on their own.

We have studied the SBDs of OpuA on the single-molecule level using optical tweezers. This has been performed in a membrane-mimicking environment by the use of nanodiscs, the protein embedded in a phospholipid bilayer surrounded by a “membrane scaffold protein” (MSP)^29^. By introducing a single-point mutation K521C, both SBDs of the homodimer are probed simultaneously in pulling experiments to uncover unfolding transitions and potential interactions. To assist in data interpretation, a single SBD was isolated (also referred to as OpuAC) to compare its unfolding pattern with the full-length membrane transporter, both experimentally and by molecular dynamics simulations. By varying the ionic strength, interactions between the SBDs (as previously observed by smFRET) were probed and the subsequent unfolding patterns were observed.

## Results

### Unfolding of both substrate-binding domains from OpuA

The transmembrane ABC-transporter OpuA from *Lactococcus lactis* has been successfully expressed and purified as described previously. The protein retains its substrate-dependent ATPase activity and ionic strength-dependent activation when reconstituted in lipid nanodiscs^21,24,28^. Therefore, OpuA is an ideal candidate for single-molecule studies on the impact of substrate and ionic strength on the conformational states and dynamics of a membrane protein. OpuA is a heterotetramer (two times the two subunits OpuAA and OpuABC), consequently a single-cysteine mutation will be repeated on the second subunit. One cysteine mutation therefore suffices to attach two handles for mechanical pulling experiments by optical tweezers.

The mutation is introduced at the back of the substrate-binding domain (SBD) near the C-terminus at position K521C, whereas the SBD is attached by a linker to the transmembrane domain (TMD) via the N-terminus of the SBD (**Figure 1B**). By pulling on residue K521C, observed length changes are expected to originate from the unfolding of both SBDs (between residue 319 and 521) and potential interactions between the SBDs.

OpuA-K521C has been purified and reconstituted in MSP1E3D1 lipid nanodiscs formed from a synthetic lipid mixture consisting of a weight ratio of 38:12:50 DOPE:DOPG:DOPC. It has previously been shown that the salt-, glycine betaine-, ATP- and c-di-AMP-dependent ATPase activities of the mutant are similar to those of the wild-type^28^.

To prevent damage of the protein by the laser light, the protein is spatially separated from the focus of the laser beam by the attachment of double stranded (ds) DNA handles. This was achieved by covalently labelling the protein with maleimide-oligonucleotides. Shortly before a measurement, the protein-oligonucleotide construct was incubated with 185-nm-long dsDNA handles. The handles form an overhang with the oligonucleotide on one side and are functionalized with either biotin or anti-digoxigenin on the other side (**Materials and Methods**)

Optical tweezer experiments were performed using a microfluidic set-up with two optical traps. First, a bead coated with streptavidin was caught in one trap. Then, a second bead coated with anti-digoxigenin, which was incubated with the protein, was caught in the other trap. Spontaneous tether formation between the two beads was initiated, through the formation of the streptavidin-biotin bond, by moving the two beads into contact with each other. The formation of a single tether could be confirmed by comparing the force response to that of a well-defined single dsDNA tether (defined by assessing whether the curve can be fitted by the extensible worm-like chain (eWLC) model, see **Materials and Methods**) when moving one bead away from the other.

To study the unfolding and folding, force-distance (FD) curves were collected. To obtain FD curves, one bead remains static and the other bead was moved at a constant velocity of 20 nm/s (unless stated otherwise). Here, the FD curve starts at a position where the protein-construct was not under tension. Multiple cycles per molecule were recorded with a waiting step of 1 s at every start- and endpoint of the curve. Unfolding of domains is observed by an abrupt discontinuity or rip in the FD curve^1^. Length changes in the protein domains were measured by fitting the FD curve with a worm-like chain (WLC) model.

Distinct unfolding steps are visible when pulling on the full transmembrane complex (**Figure 2A**). The total unfolding of both SBDs was observed, corresponding to a total length change of *∼* 150 nm. This is twice the length of 72 nm that is expected from one SBD unfolding (**Materials and Methods**). It should be noted that the total unfolding is a rare event and was only observed at a slow pulling rate of 10 nm/s; the slower rate of force application leads to a higher probability of observing the entire structure unfolding at a given applied force^30,31^.

**Figure 2.**
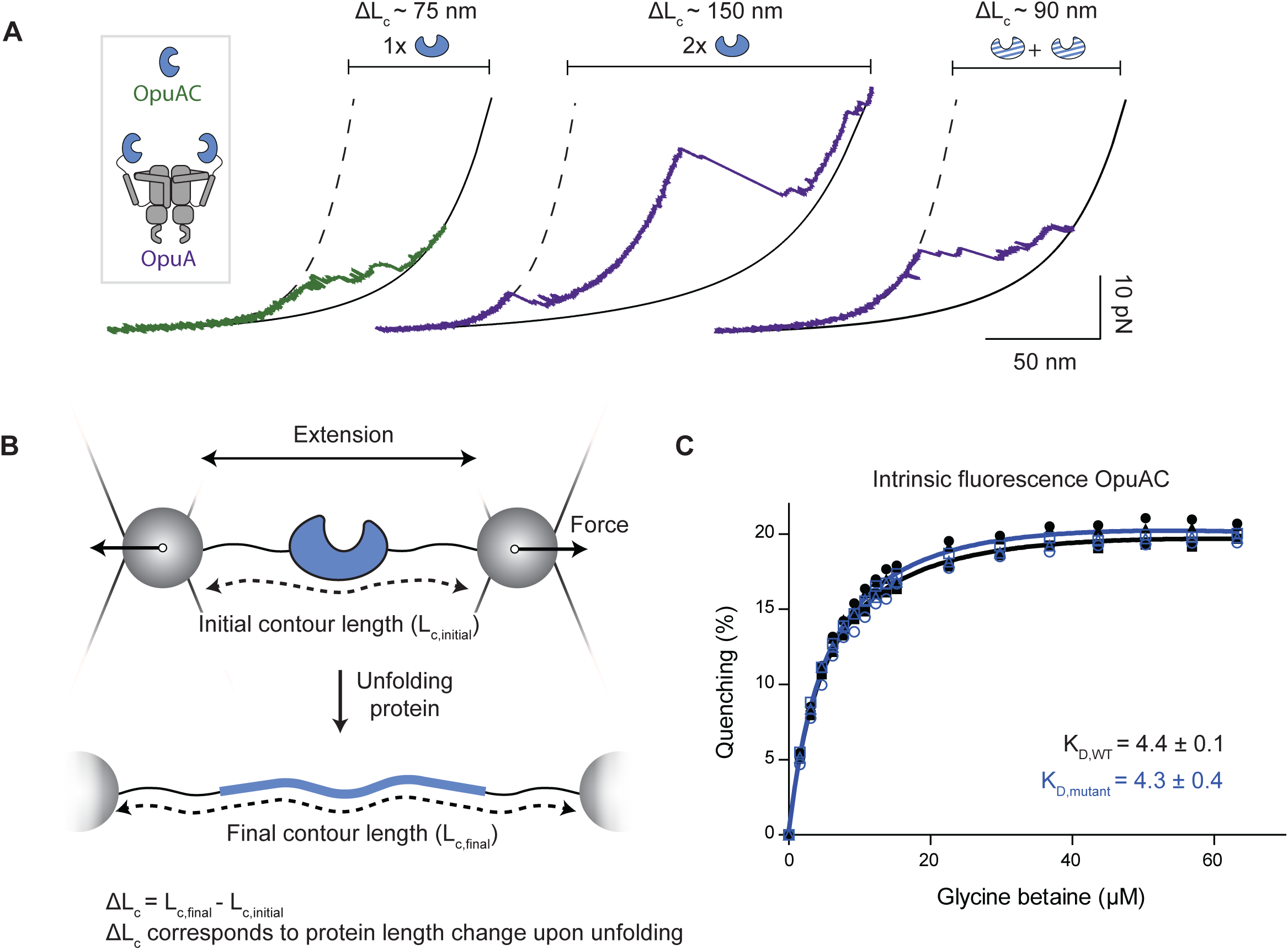
Unfolding of OpuAC and OpuA determined by contour length changes and intrinsic fluorescence of OpuAC. (**A**) Representative unfolding curves of OpuAC (green) and OpuA (purple) in 50 mM HEPES-K pH 7.0, 600 mM KCl recorded at 20, 10 and 20 nm/s from left to right. The dashed curve represents an eWLC-fit of the dsDNA linkers (protein fully folded) and the black curve the WLC-fit of the unfolded protein. Numbers above the graph represent the measured contour length change (ΔL_c_). Left: the total unfolded length of OpuAC (isolated SBD) is similar to the predicted value based on the structure (72 nm, PDB: 3L6G). Middle: at a low pulling velocity, both SBDs of OpuA can be unfolded based on the total unfolded length corresponding to two SBDs. Right: most commonly observed unfolding pattern of OpuA. Both SBDs are involved in unfolding as the contour length corresponds to more than expected from one SBD, but the SBDs do not fully unfold. (**B**) Calculation of the contour length gain. The measured change in contour length by fitting the worm-like chain model corresponds to the protein length change upon unfolding. This length change can be calculated from the structure by subtracting the final contour length (bottom) from the initial contour length (top). The final contour length can be calculated by multiplying the number of amino acids in the domain by the average length of the backbone of a single amino acid (0.365 nm) and the initial contour length can be obtained from the structure. (**C**) Intrinsic protein fluorescence-based ligand affinity measurements of wild-type OpuAC (black, closed symbols) and mutant S319C K521C OpuAC (blue, open symbols) in 20 mM HEPES-K pH 7.0, 300 mM KCl. Each symbol represents a different replicate. The colored line represents the non-linear least squares fit of Equation 1. The average and standard deviation of the K_D_ have been reported.

The most commonly observed pattern is where a total unfolded length change of *∼* 90 nm, which was frequently observed prior to breaking of a bond in between the two beads. This length change corresponds to a partial unfolding of the two SBDs (**Figure 2A**).

It could be argued that the total length change does not only include the unfolding of the SBDs, but may also involve a structural change in the TMDs. To asses this scenario, we have measured isolated soluble SBDs, named OpuAC. This construct has been examined to investigate whether the observed unfolding steps from the full-length OpuA, suggested to originate from the SBDs, are similar to the steps observed in the isolated SBD.

### Unfolding the isolated substrate-binding domain of OpuA

OpuA is a homodimer and each observed unfolding step in our FD curves could originate from either of the SBDs (or other structural elements). It is therefore challenging to assign in which order each structural element unfolds and predict the corresponding mechanical intermediates. To assist in the assignment of the unfolding steps observed, the SBD on its own (i.e., OpuAC, a soluble protein) was expressed, purified and labelled. Cysteine-mutations were introduced at the positions equivalent to S319C and K521C in the full-length protein; the first corresponds to the first residue after the membrane anchor sequence of OpuA. With these sites for labeling, we mimic the pulling geometry of the full complex (**Figure 1B**).

To confirm that the cysteine-mutations did not introduce structural changes, the K_D_ was determined by measuring changes in intrinsic tryptophan fluorescence upon binding of the substrate glycine betaine. The double mutant did not show any difference compared to wild-type OpuAC, and the dissociation constants (K_d_) were 4.3 ± 0.4 μM and 4.4 ± 0.1 μM, respectively (± standard deviation, **Figure 2C**). These values are similar to those previously reported^22^.

OpuAC was subjected to similar experimental conditions as OpuA to study the unfolding pattern by obtaining FD curves. The observed contour length change when completely unfolding OpuAC from a natively folded state of 68.1 ± 4.4 nm (N = 12, SD) matches the predicted length of 71.8 nm based on the crystal structure (PDB: 3L6G).

From the obtained FD curves, we observe four unfolding intermediates (**Figure 3**, **Table 1**). OpuAC consists of two distinct lobes. The amino acids at the C- and N-terminus are assigned to lobe 1, and lobe 2 consists of the amino acids in the middle of the chain. Due to this complex architecture, it is not trivial to assign unfolding intermediates to a structural element solely based on a crystal structure. To assist in the assignment of the unfolding pattern, steered all-atom molecular dynamics (MD) simulations have been performed to uncover the relative mechanical stability of each structural element upon the application of a pulling force (**Figure 3BC**, **Figure S1**).

**Figure 3.**
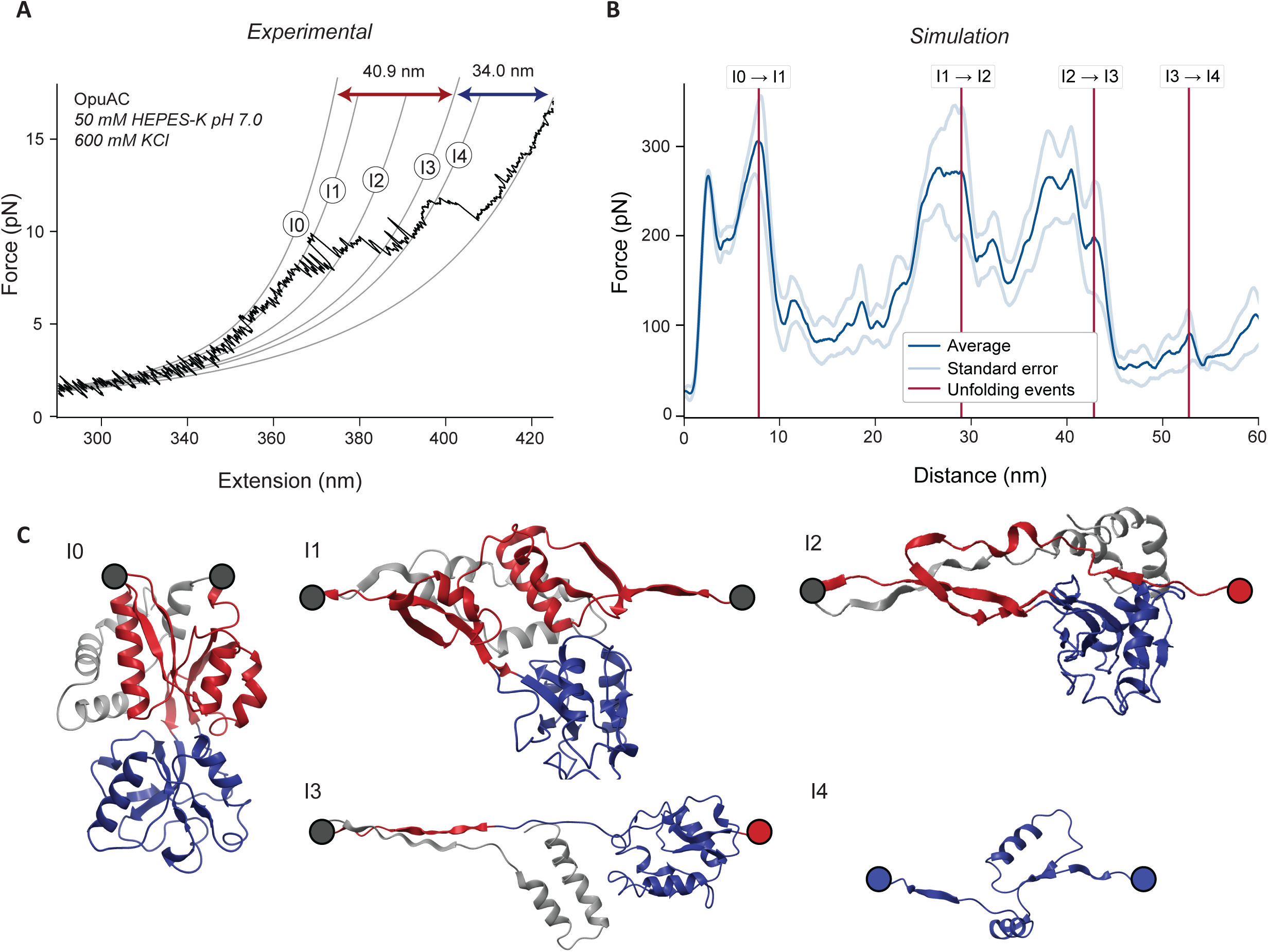
Experimental and simulated unfolding of OpuAC. (**A**) Representative experimental force-distance curve of the unfolding of OpuAC (isolated SBD). Contour length gains of the WLC-fits from the folded state (I0) to the four intermediates (I1-I4) correspond to the lengths found by MD simulations. The lengths corresponding to the colored arrows are in agreement with the expected lengths for unfolding lobe 1 (red) and lobe 2 (blue). (**B**) Average FD curve of the MD simulations, as a function of the distance between positions S319C and K521C, shown together with its standard error which is represented by the light blue lines. The contour length changes are indicated on the simulated FD curve by the vertical lines. See also **Figure S1**. (**C**) Structures of the mechanical intermediate states found by MD simulations based on the crystal structure (PDB: 3L6G). Colors in the structure represent lobe 1 (red), lobe 2 (blue) and the C-terminal tail (light gray). The drawn circles represent the attachment point of the handle (dark gray) or an unfolded amino acid chain corresponding to (part of) lobe 1 (red) and/or lobe 2 (blue), which has been cropped for simplicity of the figure.

**Table 1:** Calculated, simulated and measured contour length changes when unfolding OpuAC, data correspond to measurements shown in Figure 3. Expected contour length calculated as described in Materials and Methods where folded ➔ I1 represents unfolding of the N-terminal beta sheet, I1 ➔ I3 unfolding of lobe 1 and I3 ➔ unfolded unfolding of lobe 2 to the fully unfolded state. Length changes in the MD simulations correspond to the distance between residue S319C and K521C with intermediates as shown in **Figure 3**.

|  | Expected contour length change (nm) | Length change MD simulations (nm) | Measured contour length change (nm) | Unfolding force (pN) |
| --- | --- | --- | --- | --- |
| **Folded → I1** | 6 (6) | 8 (8) | 8 (8) | 5.9 |
| **I1 → I3** | 29 (35) | 34 (42) | 33 (40) |  |
| **I1 → I2** |  | 20 (28) | 17 (25) | 9.8 |
| **I2 → I3** |  | 14 (42) | 16 (41) | 9.5 |
| **I3 → Unfolded** | 37 (72) | 32 (74) | 34 (75) |  |
| **I3 → I4** |  | 9 (51) | 9 (50) | 10.5 |
| **I4 → Unfolded** |  | 23 (74) | 25 (75) | 11.6 |

Based on the all-atom simulations, the following order of unfolding can be assigned: first, the N-terminal beta sheet from lobe 1 starts to unfold. Then, the remaining structure of lobe 1 unfolds and finally lobe 2 unravels (**Figure 3C**). The expected contour length changes based on the crystal structure and MD simulations match the experimentally obtained lengths (**Table 1**). We have therefore assigned the unfolding intermediates from the experimental FD curves similarly as predicted from MD simulations, where first lobe 1, and then lobe 2 unfolds (**Figure 3**). Note that the unfolding forces of the MD simulations are a magnitude larger than the experimental values. This can be explained by the high pulling velocity used in the simulations compared to the experiments (0.1 m/s *versus* 20 nm/s, see also **Materials and Methods**)

Notably, two out of seven replicates revealed that also transient beta sheets could be formed within the intermediate structure where the unfolded amino chain could form a β-sheet with the N-terminal part of the protein. These short-lived beta sheets result in a significantly higher unfolding force contributing to a larger error in the unfolding force. Nonetheless, the order of unfolding remained similar to the other replicates.

### Glycine betaine mechanically stabilizes OpuAC

Ligands can stabilize their corresponding binding partners, which can for instance be observed by an increase in the thermostability of the protein. In single-molecule force experiments, this effect can be observed as an increase in unfolding force and/or a change in the unfolding pathway^32–34^. Glycine betaine binds OpuAC with a K_D_ of 4.4 μM (**Figure 2C**). Therefore, we have performed similar optical tweezers experiments on OpuAC also in the presence of 100 μM glycine betaine.

From the obtained FD curves, we observe a similar total unfolding length of 71.0 ± 4.5 nm (N = 12, SD) to the apo conditions. Additionally, similar unfolding intermediates are observed as in the apo state (based on the observed length changes), though the forces of each unfolding step are significantly higher in the presence of glycine betaine, indicating a mechanical stabilizing effect of the ligand (**Figure 4**).

**Figure 4.**
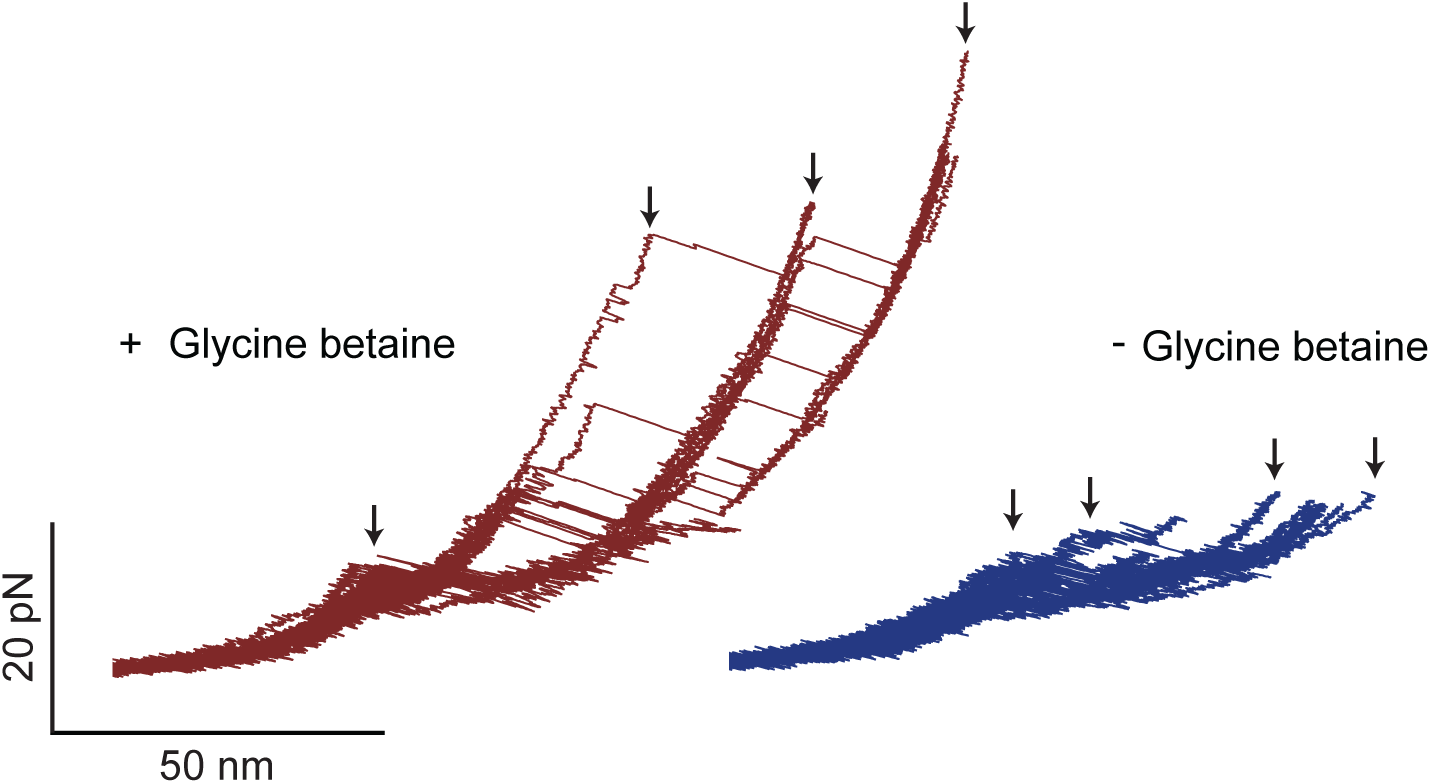
Mechanical stabilization by the substrate glycine betaine of OpuAC. Overlay of the unfolding traces in the presence of 100 μM glycine betaine (left, 4 molecules, 33 traces) and in the absence of glycine betaine (right, 9 molecules, 22 traces) in 50 mM HEPES-K pH 7.0, 600 mM KCl. Traces with the substrate present show higher unfolding forces, indicating mechanical stabilization by the ligand. Stars represent similar unfolding intermediates in both conditions based on contour length changes.

### The SBD shows similar unfolding steps in the full complex of OpuA

After determining the steps involved in the unfolding signature of an isolated SBD in OpuAC, FD curves of the full-length transmembrane transporter OpuA were performed in 50 mM HEPES-K pH 7.0, 600 mM KCl. The unfolding pattern in the FD curves of OpuA showed similar sized, reversible unfolding steps as the isolated SBD OpuAC (**Figure 5**, **Table S1**). Correspondingly, unfolding occurs at a comparable force range in both constructs. Occasionally, we observed a non-unfolding intermediate, which later started to unfold again when recording consecutive FD curves on a single molecule (**Figure S2**)

**Figure 5.**
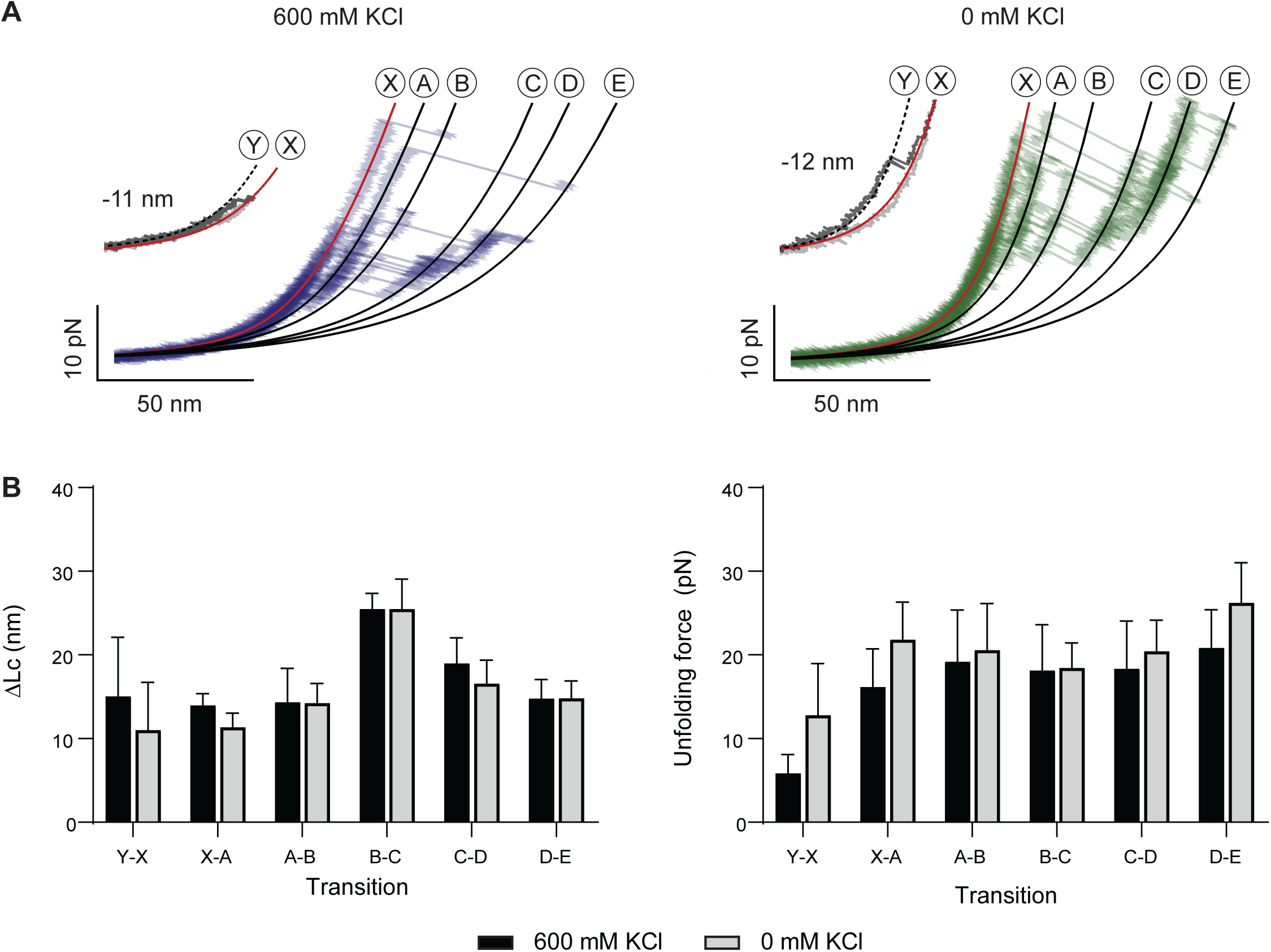
Unfolding of OpuA with and without additional salt. (**A**) Overlay force-distance curves of consecutive unfolding traces of a single molecule measured in a buffer of 50 mM HEPES-K pH 7.0, with (left) and without (right) additional 600 mM KCl. Intermediates with WLC-fits have been labeled according to the order of occurrence. The gray inset represents the first pulling cycle, whereas the red line in the figure and inset correspond to the same length. (**B**) Distribution of the contour length increase (left) and unfolding force (right) represented by a bar plot. Data points have been collected from 6 molecules, 28 traces (600 mM KCl) and 4 molecules, 34 traces (0 mM KCl). Error bars represent the standard deviation, see also **Table S1 and Figure S3**.

### Interactions between SBDs probed at different ionic strengths

SmFRET results have suggested an interaction between the SBDs, of which the state where the SBDs are in close proximity is more prominent in low salt conditions (<100 mM KCl in 50 mM HEPES-K, pH 7.0)^28^. Therefore, the full-length transmembrane protein OpuA was subjected to force in 50 mM HEPES-K pH 7.0 without additional KCl. The unfolding pattern of OpuA is similar in both salt conditions (**Figure 5A**), with unfolding steps of similar length and comparable unfolding forces (**Figure 5B**).

Notably at low salt, an additional event is observed prior to the reversible unfolding pattern (inset **Figure 5A**), where the SBDs move from being at close proximity to being further away from each other. This event occurs almost exclusively at the start of a measurement, when the protein has not yet been subjected to a pulling force. Only rarely can this event be observed again after unfolding the protein, in contrast to the reversible folding and unfolding of the SBDs (e.g., observed in *∼* 20% of the traces in low salt condition *versus ∼* 90% for the subsequent unfolding step in case of 600 mM KCl, N = 28). As this event cannot be assigned to an unfolding step based on the unfolding pattern of OpuAC and due to its occurrence prior to any other unfolding events, we have assigned this event to an interaction between the SBDs (**Table S1**).

As noted above, the population of the SBDs in close proximity is higher in no salt conditions compared to high salt. Therefore, in the absence of salt, we expect in our optical tweezer measurements a higher fraction of molecules where this interaction is observed and/or a higher required force to break this interaction.

We observe a contour length change of this interaction of 11 ± 5 nm and 15 ± 6 nm (**Figure S3**, **Table S1**) in 50 mM HEPES-K pH 7.0, with and without 600 mM KCl, respectively. While both length changes are in the same order of magnitude, a relatively large error is observed which can be attributed to the non-linearity of the WLC fit in the lower force region and changes in stiffness of DNA by varying salt concentrations. Notably, the force required to break this interaction is higher in a buffer without additional salt. This is in line with the observed FRET distributions at low and high salt conditions^28^.

## Discussion

We have used optical tweezers to examine the unfolding and potential interactions of the SBDs from the membrane protein OpuA on the single-molecule level. Additionally, the isolated SBD (also referred to as OpuAC) was studied, as a reference for the mechanical unfolding signature.

Unfolding OpuAC showed similarly sized unfolding steps as in the full-length transporter-complex. Steered all-atom MD simulations aided in the interpretation of the order of unfolding of the secondary structure elements in the isolated SBD revealing several mechanical intermediates. Each of the simulations was consistent in assigning the least mechanical stable structural element (N-terminal β-sheet) and the order of unfolding of each lobe. One of the simulations revealed that also the unfolded amino chain could form a secondary structure (β-sheet) with the C-terminal part of the protein, resulting in a significantly higher unfolding force. This may explain the observation of occasional “non-unfolding”/misfolded states, where no unfolding steps were observed under force (**Figure S2**).

The substrate glycine betaine mechanically stabilizes the SBD, which was observed as an increase in the unfolding force (with exception of the first unfolding step, *vide infra*). Substrate binding has been shown to stabilize other proteins^33,35^. Comparing the positions of the amino acids involved in the binding of glycine betaine (**Figure S4**, PDB: 3L6H) to the intermediates from the MD simulations, hints as to why the first unfolding step is not stabilized. Most amino acids from the binding pocket are only involved in unfolding in the later mechanical intermediates. Another hypothesis of the elevated unfolding forces could be the general stabilization of the protein by the substrate without being specifically bound to the identified substrate-binding site, as glycine betaine is known to be a stabilizing osmolyte. However, this effect has only been observed in studies working at a concentration of >100 mM^36,37^, while we have observed stabilization at 0.1 mM.

The isolated SBD and the full-length transporter showed similar unfolding steps, supporting the assumption that the observed length changes originate from only the SBDs. In case of the full-length membrane transporter OpuA, we have studied the unfolding of the SBDs with and without the addition of salt. Both conditions showed a similar unfolding pattern with similar length changes. Though in both cases, we rarely observed the complete unfolding of both SBDs where one (or a part of both SBDs) is shielded from unfolding. It could be argued that this might be caused by the interference of the charged lipids and/or negatively charged nanodisc. However, we also observed this partial unfolding in the presence of 600 mM KCl, which would shield some of these electrostatic interactions. Another possibility would be the formation of a non-native, mechanically stable secondary structure during the unfolding of the protein, similar to what has been shown with the MD simulations of the isolated SBD OpuAC.

Our data on OpuA also suggests that there is an interaction of the SBDs prior to unfolding. The force required to break this interaction was higher at low salt (buffer only) than in the presence of buffer plus 600 mM KCl. This increase in force matches the observation of a larger high FRET population in the low salt condition from smFRET data^28^. Additionally, the experimental values for the change in length 11 ± 5 nm and 15 ± 6 nm in buffer without and with salt, respectively, are in the same order of magnitude as expected from an interaction (**Materials and Methods, Figure 6**).

**Figure 6.**
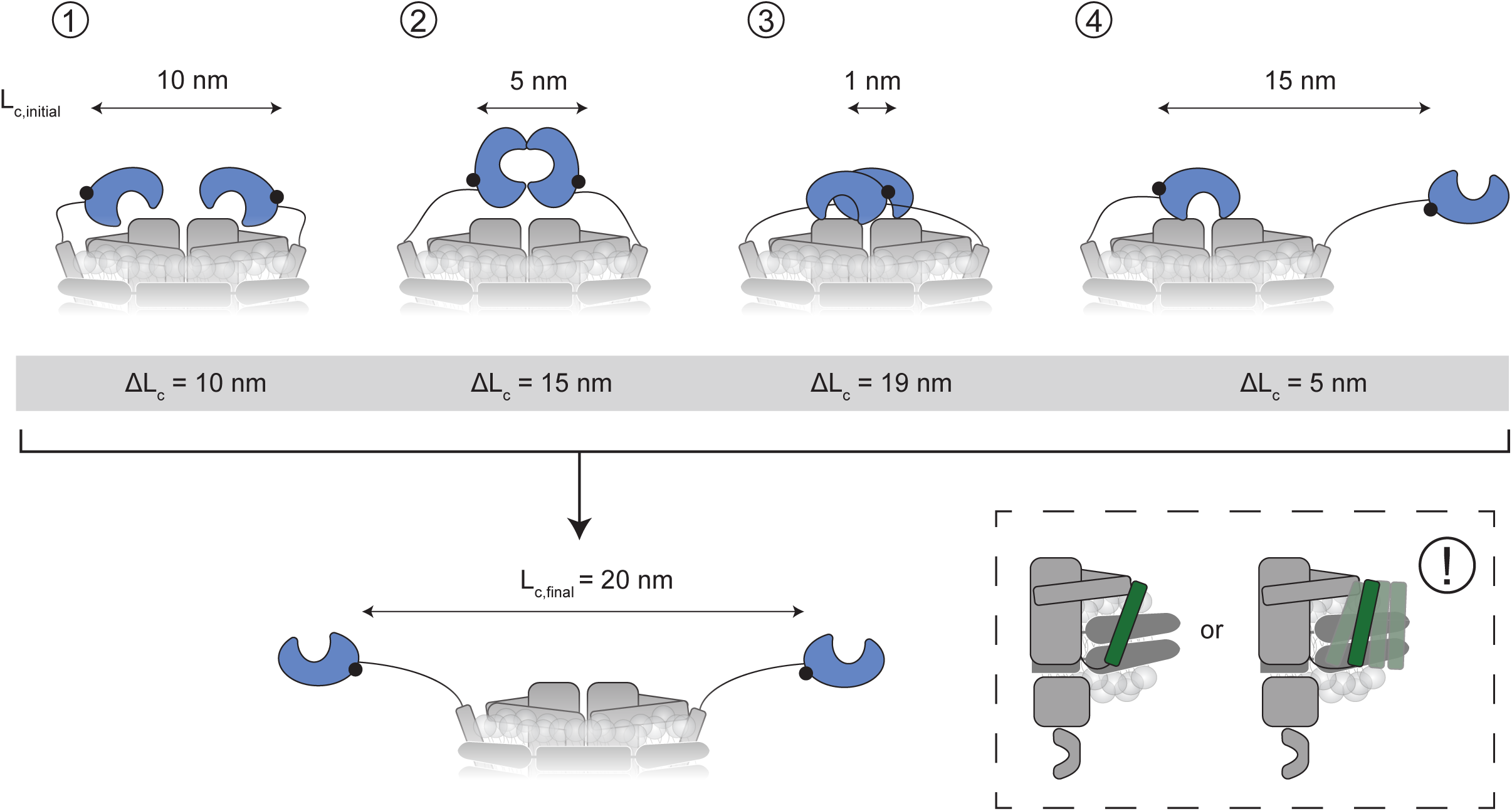
Predicted contour length changes based on different interactions of the SBDs. Four different scenarios of different conformations of the SBDs, shown with their initial distance (L_c,initial_, top) and the expected contour length change (ΔL_c_) calculated based on the extended length (L_c,final_, bottom) of 20 nm. Note that the numbers shown in the figure are approximations and possible flexibility of the anchoring helix has not been taken into account (bottom right, see also **Materials and Methods).** (**1**) Interaction with the TMD as resolved in the low-salt cryo-EM structure as reported by van den Noort et al.^28^, (**2**) the lobes interact on the side of the substrate-binding pocket, (**3**) each SBD interacts with the transmembrane domain where the maximum distance has been calculated based on the length of the linkers, and (**4**) docking of one of the SBDs where the second SBD is parallel to the pulling direction.

There are several scenarios in which the two SBDs could be in close proximity. From these scenarios, expected length changes can be calculated. These scenarios include different interactions and corresponding positions of the SBDs: either the SBDs interact with each other, the SBDs interact with another part of the protein or one of the SBDs is docked (as illustrated in **Figure 6**). Although a low-resolution cryo-EM structure in low salt is available, some approximations have to be made when calculating the contour length changes, resulting in a larger uncertainty in the calculated values.

The first source of uncertainty is the low resolution of the SBDs in the low-salt structure and the unknown relative orientation of the SBDs in the scenarios where they interact. Extreme cases have been taken into account in **Figure 6**. Hence, the reality may be a structure that is in-between one of the proposed scenarios.

Second, the possibility of the movement of the anchoring helix in the lipid nanodisc increases the uncertainty. This can depend on how tightly the lipids are packed and how the charged lipids behave at different concentrations of salt under the application of force on the protein. Based on earlier data, the anchoring helix is more flexible in high salt conditions. The anchoring helix is under high salt conditions more likely to move towards the edge of the nanodiscs, potentially adding extra length. This may explain the differences in experimental values for the length change in the absence and presence of 600 mM KCl (11 ± 5 nm and 15 ± 6 nm, respectively).

On the basis of the aforementioned approximations in expected contour length and the observed length changes, docking (scenario 4) is the least likely. An interaction with the TMD (scenario 1) or between both SBDs (scenario 3) are the most probable scenarios for the interaction of the SBD (**Figure 6**).

The observation of a potential interaction of the two SBDs raises the question of why proteins would have differences in domain-domain interactions at different salt concentrations. In the context of OpuA, interacting SBDs may serve as a mechanism to facilitate cooperativity of the two SBDs to enhance transport by retaining the SBDs in near proximity.

In conclusion: we show that optical tweezers can be used to study the conformational dynamics of integral membrane proteins under native-like and physiologically relevant conditions. Information that can be obtained includes not only the force that is associated with an event but also the distance between the two attachment points. We observe weak interactions between the two substrate-binding domains, and we determine their unfolding trajectories. Using a nanodisc as a membrane-mimicking environment for the ABC transporter OpuA offers the opportunity to also study dynamics on other parts of the protein using optical tweezers. The methods presented in this work provide an opportunity to start addressing many outstanding questions relating to how a membrane protein functions and (un)folds in a native-like membrane.

## Acknowledgements

We would like to thank M. van den Noort, P. Drougkas and V.B. Verduijn for critical reading of the manuscript. We acknowledge funding from the Dutch Research Council: NWO OCENW.KLEIN.526.

## Author contributions

L.S. conducted the experiments and analyzed the data, and wrote the manuscript with the help of K.T., J.S. performed the simulations and wrote the corresponding section of the manuscript. L.S., J.S., B.P. and K.T. interpreted experiments. All authors supervised and revised the manuscript.

## Materials and Methods

### Expression and purification of MSP1E3D1

MSP1E3D1 was expressed and purified with slight modifications from a previously described method^38^.

#### Large scale expression of MSP1E3D1

*E. coli* BL21(DE3) cells with the pMSP1E3D1 plasmid were grown aerobically in 2 L fermenters at 37 °C in phosphate buffered TB medium (Formedium, UK) supplemented with antifoam and 10 μg/mL Kanamycin. The cells were induced with 1 mM IPTG (isopropyl 1-thio-β-D-galactopyranoside) at an OD_600_ of 2.0 and were allowed to grow for 3 more hours. Subsequently, the cells were harvested by centrifugation (6,000 g, 15 min, 4 °C), washed with 50 mM KPi pH 7.8, resuspended in 200 mL 50 mM KPi pH 7.8 and finally flash-frozen in liquid nitrogen to be stored at - 80 °C.

#### Purification and His-tag cleavage

For purification of the MSP1E3D1 protein, 100 μg/mL DNase, 1 mM PMSF and 1% TritonX-100 were added to 100 mL of cells while stirring. The cells were broken by sonication (5 sec on, 5 sec off, 70% amplitude), centrifuged (30,000 g, 30 min, 4 °C) and the supernatant was supplemented with 20 mM Imidazole pH 8.0. MSP1E3D1 was purified using affinity chromatography. The protein was allowed to bind to washed Ni^2+^-Sepharose (7.5 mL column volume) for 60 min at 4 °C while gently agitating. Subsequently, the column was washed first with 5 CV Buffer A (50 mM Tris-HCl pH 8.0, 0.3 M NaCl) supplemented with 1% Triton X-100, secondly with 5 CV Buffer A supplemented with 50 mM Na-cholate and 20 mM Imidazole, and finally with 5 CV Buffer A supplemented with 50 mM Imidazole. The protein was eluted in fractions of 2 mL with Buffer A supplemented with 500 mM Imidazole. The fractions containing protein were combined.

The His-tag was cleaved by incubation with TEV protease (40:1 w/w ratio) and 5 mM EDTA for 90 min at 4 °C while gently agitating. The buffer was exchanged by dialysis overnight at 4 °C against 1 L dialysis buffer (50 mM Tris/HCl pH 7.4, 0.1 M NaCl, 0.5 mM EDTA, 0.5 mM DTT) in boiled Servapor® dialysis tubing with a diameter of 16 mm. Further protein purification was achieved with affinity chromatography using Ni^2+^-Sepharose (5 mL CV). The protein solution was first incubated with washed Ni^2+^-Sepharose for 30 min at 4 °C while gently agitating. Then, the column material was drained, and the flow through was collected. Subsequently, the column was washed with 4 CV wash buffer (20 mM Tris/HCl pH 7.4, 0.1 M NaCl, 25 mM imidazole) of which 2 mL fractions were collected. The fractions containing protein were combined and concentrated to 5 mg/mL using a Vivaspin 20 ultrafiltration device with a molecular weight cut-off of 10 kDa. Finally, aliquots were stored at −80 °C after flash freezing in liquid nitrogen.

### Large scale expression of OpuA or OpuAC

*L. lactis* Opu401 cells carrying pNZOpuAHis or pNZOpuACHis were grown in 2% (w/v) Gistex, 1% (w/v) glucose, 65 mM NaPi pH 7.0 plus 5 μg/ml chloramphenicol at 30 °C under semi-anaerobic conditions using 2 L pH- and temperature-controlled fermenters while stirring at 200 rpm. The pH was kept at 6.5 using 4 M potassium hydroxide. Cells were induced with 0.05% (v/v) of a nisin-producing strain at an OD_600_ of 2.0, then supplemented with additional glucose (1% (w/v)), and the cells were grown for 2 more hours. Subsequently, the cells were harvested by centrifugation (6,000 g, 15 min, 4 °C), washed with 100 mM KPi pH 7.5 and resuspended in 50 mM KPi pH 7.5 (supplemented with 20% (v/v) glycerol in case of OpuA) to a final OD_600_ of *∼* 200. The cells were flash-frozen in liquid nitrogen to be stored at −80 °C.

### Purification and reconstitution of OpuA

OpuA was purified and reconstituted as described before, for a more detailed protocol see^28^.

#### Preparation of OpuA Crude Membranes

The cells with an OD_600_ of *∼* 200 were thawed, 100 μg/mL DNAse and 2 mM MgSO_4_ were added and the cells were broken by passing them twice through a high pressure device (maximator; 30 kPsi). Afterwards, 5 mM EDTA pH 8.0 and 1 mM PMSF were slowly added while stirring. Cell debris was removed by low-speed centrifugation (15,000 g, 15 min, 4 °C). Subsequently, the supernatant was spun down at 184,842 g for 75 min at 4 °C. The pellet was washed in 25 – 50% of the original volume using 50 mM KPi pH 7.5 plus 20% (v/v) glycerol. The washed pellet was resuspended in 50 mM KPi pH 7.5 plus 20% (v/v) glycerol (*∼* 12 mL per 2 L culture) and stored at −80 °C after being flash-frozen in liquid nitrogen. Protein concentrations were determined using the Pierce BCA Protein Assay Kit.

#### Purification, reconstitution and labelling of OpuA in Nanodiscs

Crude membranes containing OpuA were solubilized in 200 mM KCl, 50 mM KPi pH 7.0, 20% (v/v) glycerol and 0.5% (w/v) DDM with a final concentration of 3 mg/mL membrane protein in the solubilization mixture. The mixture was incubated for 30 min on ice and mixed once by pipetting up and down after 15 min. Insolubilized membranes were spun down (336,896 g, 20 min, 4 °C). The solubilized protein was added to a closed column containing washed Ni^2+^-Sepharose (0.5 mL CV) in the presence of 10 mM imidazole and was allowed to bind while gently pipetting up and down. The resin was left to sediment and the outlet of the column was opened. The resin was washed using 20 CV of buffer B (200 mM KCl, 50 mM KPi pH 7.0, 20% (v/v) glycerol) supplemented with 0.04% (w/v) DDM and 50 mM imidazole). OpuA was eluted in one fraction of 0.6 CV and five subsequent fractions of 0.4 CV Buffer B plus 0.04% DDM and 200 mM imidazole.

For reconstitution, lipids were prepared with a ratio of 50 mole percent (mol %) 1,2-dioleoyl-sn-glycero-3-phosphoethanolamine (DOPE), 38 mol % 1,2-dioleoyl-sn-glycero-3-phospho-(111-rac-glycerol) (DOPG), and 12 mol % 1,2-dioleoyl-sn-glycero-3-phosphocholine (DOPC). 6.25 mg/mL lipids were tip sonicated in an ice-water bath for 8 cycles (15 sec on, 45 sec off at an intensity of 4 μm) and subsequently solubilized in 1% (w/v) DDM. The reconstitution mix consisted of 900 μM lipids, 4.5 μM OpuA (tetrameric complex), 45 μM MSP1E3D1 and 50 mM KPi pH 7.0 in a final volume of 700 – 900 μL. After nutation for 1 h at 4 °C, 500 mg SM-2 BioBeads were added and the mixture was incubated overnight at 4 °C under gentle agitation. The beads were removed by transferring the solution to a clean Eppendorf tube using a needle. Subsequently, the aggregates were removed by centrifugation (20,817 g, 15 min, 4 °C). In case of cysteine mutants of OpuA, 1 mM DTT was supplied during the breaking of the cells, purification and reconstitution.

For removal of the empty nanodiscs, the protein after reconstitution was loaded to a column containing washed Ni^2+^-Sepharose (0.2 mL CV). The flow-through was collected and reapplied to the column twice.

In case of non-labelled OpuA, the column was washed with 10 CV buffer C (300 mM KCl, 20 mM HEPES-K pH 7.0) supplemented with 25 mM Imidazole. The protein was eluted with 3 CV buffer C supplemented with 200 mM imidazole. In case of a non-labelled cysteine mutant, 1 mM DTT was also added to buffer C.

For labelled OpuA, the column was washed with 5 CV buffer C. The column was closed, the maleimide-oligonucleotides were added in excess and the protein was incubated on ice for 4 h without nutation while wrapped in foil. Then column, was washed with 10 CV buffer C supplemented with 25 mM Imidazole. The protein was eluted with 2 CV buffer C supplemented with 200 mM imidazole.

The OpuA-nanodiscs were further purified by size-exclusion chromatography using a Superdex 200 Increase 10/300 GL column in buffer C (supplemented with 1 mM DTT in case of non-labelled cysteine mutants).

### ATPase assay of OpuA in nanodiscs

The ATPase activity of OpuA in nanodiscs was determined with a two-enzyme coupled enzyme assay as described by Akira et al.^24^. The assay was performed in technical triplicates, in a 96 well-plate. Each replicate consisted of 12 different concentrations of one of the four variables: Mg-ATP, KCl, glycine betaine or cyclic-di-AMP. A standard solution of 200 μL per well was composed of 50 mM HEPES-K (pH 7.0), 4 mM phopho(enol)pyruvic acid, 600 μM β-nicotinamide adenine dinucleotide, 4.5 μL pyruvate kinase/lactic dehydrogenase enzyme mixture from rabbit muscle (Sigma-P0294-5 mL) with 600 – 1000 units PK/mL and 900 – 1400 units LDH/mL, 450 mM KCl, 100 μM glycine betaine, 10 mM Mg-ATP and 0 mM cyclic-di-AMP. The samples were incubated for 5 minutes at 35 °C. Subsequently, either 10 mM Mg-ATP or 100 μM glycine betaine was titrated to each well and the solution was briefly mixed. The absorbance of NADH at 340 nm was monitored for 25 min with the Tecan Spark multimode reader. The absorbance was corrected for path length using the absorbance at 900 and 977 nm of each well, measured at the end of each experiment.

### Isolation and purification OpuAC

#### Isolation of cytosolic protein OpuAC

Cells with an OD_600_ of *∼* 200 were thawed, 100 μg/mL DNAse and 2 mM MgSO_4_ were added and the cells were broken by passing them twice through a high-pressure device (maximator; 30 kPsi). Afterwards, 5 mM EDTA pH 8.0 and 1 mM PMSF were slowly added while stirring. The cytosolic fraction was isolated as supernatant by high-speed centrifugation (125,171 g, 90 min, 4 °C) and stored in aliquots at −80 °C after being flash-frozen in liquid nitrogen. Protein concentrations were determined using the Pierce BCA Protein Assay Kit.

#### Purification of OpuAC

Ni^2+^-Sepharose resin (0.5 mL CV) was washed with buffer D (200 mM KCl, 50 mM KPi pH 7.0) and 40 mg of cytosolic protein was added to the resin supplemented with 10 mM imidazole. The mixture was mixed by pipetting, the resin was left to sediment and the outlet of the column was opened. The resin was washed with 20 CV buffer D supplemented with 50 mM imidazole. The protein was eluted once with 0.6 CV and 4 times with 0.4 CV buffer D supplemented with 500 mM imidazole. Fractions containing protein were combined and were further purified by size-exclusion chromatography using a Superdex 200 Increase 10/300 GL column in buffer C (300 mM KCl, 20 mM HEPES-K pH 7.0). In case of cysteine mutants of OpuA, 1 mM DTT was added during the breaking of the cells and purification.

#### Labelling of OpuAC

The purified cysteine-mutant was loaded to a column containing washed Ni^2+^-Sepharose (0.1 mL CV). DTT was removed by washing the column with 10 CV buffer C. Maleimide oligonucleotides were slowly added in excess and the sample was incubated at RT for 4.5 h. Unbound maleimide oligonucleotides were removed by washing with 8 CV buffer C. The labelled protein was eluted with 4 CV buffer C supplemented with 500 mM imidazole and further purified by size-exclusion chromatography using a Superdex 200 Increase 10/300 GL column in buffer C.

### Fluorescence measurements

Fluorescence titration experiments were performed as described by Wolters et al.^22^ with a Jobin Yvon SPEX Fluorlog FL3-22 spectrophotometer at RT. To a quartz cuvette, 800 μL of 0.5 μM purified protein (in buffer C: 300 mM KCl, 20 mM HEPES-K pH 7.0) was added. Under constant stirring, a solution of 500 μM glycine betaine in buffer C was added in steps of 2 or 10 μL using a Hamilton syringe pump (Harvard apparatus). Excitation and emission wavelengths of 295 and 360 nm (and slit widths of 1 and 5 nm), respectively, were used. The signal was collected for 20 seconds after addition of glycine betaine.

The fluorescence intensity was corrected for the baseline by subtracting the signal from titration with only buffer C (without glycine betaine). The K_D_ was calculated using the formula:

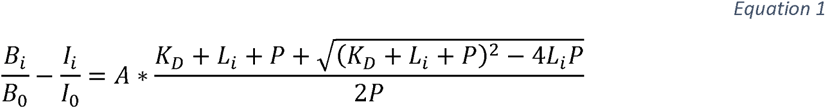

where I_i_ is the mean fluorescence intensity, B_i_ the mean intensity of the corresponding baseline and L_i_ the ligand concentration at interval i. I_0_ and B_0_ are the mean fluorescence intensities at the first interval. The protein concentration P was fixed at 0.5 µM. Both the K_D_ and A (asymptote) were defined as free fit parameters.

All measurements have been performed in triplicate. The K_D_ has been calculated for each individual experiment.

### Method details optical tweezers

For optical trapping experiments, commercially available dual-trap optical tweezers (C-trap, LUMICKS B.V., NL) were used with a microfluidic chip. Measurements were performed using a dumbbell conformation: the protein of interest was attached in between two 1 μm-sized silica beads (Spherotech, US), using double stranded (ds) DNA handles (contour length of 185 nm each) on either side of the protein. First, a short oligonucleotide (34 bases) was covalently bound to the protein via a cysteine-maleimide bond. The oligonucleotide was linked to the dsDNA handle of a length of 544 base pairs (bp) via an overhang. The 5’-end of the dsDNA handle either contained a multi-biotin or a multi-digoxigenin site to facilitate binding to the streptavidin or anti-digoxigenin beads, respectively. The protein-DNA construct was incubated on anti-digoxigenin beads prior measurements.

A typical optical tweezer measurement was performed as follows: first, a streptavidin-coated silica bead was caught in one trap. Then, an anti-digoxigenin-coated silica bead with the protein attached was caught in the other trap, making use of the multiple channels created by laminar flow in the microfluidic chip. Flow was released and the spring constants of the beads were calibrated using the thermal noise method^39^ in a channel with the appropriate buffer (50 mM HEPES pH 7.0 with or without 600 mM KCl) supplemented with an oxygen scavenger system to prevent damage of the protein by the laser (1700 U/mL catalase, 26 U/mL glucose oxidase and 0.66% (w/v) glucose). Trap stiffness of the optical traps was set to 0.3 – 0.4 pN/nm. Tether formation was probed by bringing the two beads in close contact to each other. The formation of a tether was confirmed if the force response matched that of dsDNA stretching with a contour length of 370 nm^12^. Then, the protein was subjected to multiple stretch-and-relax cycles at 20 nm/s (unless mentioned otherwise), with a 1 s waiting step at every start- and end-point of the cycle.

### Preparation of dsDNA handles

Maleimide-oligonucleotides were ordered commercially (Biomers, DE). dsDNA handles with a contour length of 185 nm were produced in house utilizing λ-phage DNA as template by PCR using Taq polymerase with the primers as listed below.

Oligonucleotides:

- Multi bio primer: 5’- Biotin-GGCGA(Biotin-dT)CTGG(Biotin-dT)CGTTGATTTG -3’
- Multi dig primer: 5’- Digoxigenin-GGCGA(Dig-dT)CTGG(Dig-dT)CGTTGATTTG -3’
- Linker primer for DNA handles: 5’- CGACTCGCTGGTCTGGTTGAACGTCAGCCCTGCC(dspacer)CCTGCCCGGC
- TCTGGACAGG -3’
- Lambda DNA: N6-methyladenine-free, N3013L
- Maleimide oligonucleotides: GGCAGGGCTGACGTTCAACCAGACCAGCGAGTCG-Maleimide

### Quantification and statistical analysis

#### Constant velocity experiments

All force-extension traces were recorded at a constant pulling velocity of 20 nm/s, unless mentioned otherwise, with a 1 sec pause at every start- and end-point of each curve. The displayed force-extension traces at 20 nm/s in the figures have either been down-sampled to a sampling rate of 7.8 kHz and filtered by a sliding average of 201 points or a sampling rate of 0.78 kHz and filtered by a sliding average of 21 points.

#### WLC fitting

The worm-like chain (WLC) model was used to describe the elastic response of the dsDNA handles and the unfolded protein. In case of dsDNA, an extensible WLC described this elastic response with a persistence length of 20 nm, temperature of 297 K and a Hookean contribution (K-value) of 400 – 1200 pN. The total contour length of the two dsDNA-handles in the dumbbell configuration of 370 nm (2 * 544 *bp*, 0.338 *nm/bp* for B-form DNA) was used. A standard WLC fit was applied in case of unfolded protein with a fixed persistence length of 0.7 nm^12^.

### Assignment of contour length gains to structure

Identification of lengths of the unfolded protein domains was performed by comparing the contour length gain obtained from the WLC-fitting to the expected length of the corresponding domain (see also **Figure 2B**). The expected change in contour length can be calculated with the following formula:

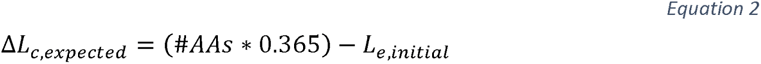

Where *#AAs* is the number of amino acids in the domain, *0.365* is the average length of the backbone of a single amino acid in nm and *L_e,initial_* is the initial distance between the N- and C-termini of the domain from the crystal or cryo-EM structure. Contour length changes that cannot be assigned to a structural element unfolding, can be attributed to a relative distance change between domains, e.g., breaking of an interaction between two domains.

Expected contour length changes from domain-domain interactions have been calculated using the low-salt model of OpuA based on cryo-EM studies^28^. Note that the substrate-binding domain has not been well resolved due to flexibility and therefore the calculated numbers serve as an estimate. Indications of the expected length changes have been made based on a nanodisc with a diameter of 11 – 12 nm.

### Molecular dynamics simulations

The molecular dynamics (MD) simulations in this study were conducted using GROMACS 2021.5^40^, with the CHARMM36 force field^41^ and the TIP3P water model. We used LINCS holonomic constraints for all bonds with hydrogen atoms^42^, this allowed us to use a time step of 2 fs in all simulations. Furthermore, we employed a Bussi−Parrinello velocity rescaling thermostat at 300 K with a time constant of 0.1 ps for both the protein and the non-protein atoms separately^43^. For the simulations that are performed under constant pressure, an isotropic Parrinello−Rahman barostat was used, set to 1 bar with a time constant of 1 ps and a compressibility of 4.5·10^−5^ bar^−1 44^. The short-ranged non-bonded interactions were evaluated using a Verlet neighbor list that was updated with a frequency tuned automatically by GROMACS. The long-range electrostatics were treated through a fourth-order particle mesh Ewald (PME) method with a real space cutoff radius of 1.25 nm and a 0.16 nm grid spacing for the FFT^45^. Before the MD simulations, all systems were submitted to an energy minimization protocol.

To start, the structure of OpuAC, with the appropriate Cysteine mutations (K521C and S319C), was generated using AlphaFold2^46^ in a ColabFold notebook^47^. The protein structure is pre-equilibrated before starting the steered MD (SMD) simulation. As part of the equilibration protocol, the protein was solvated and neutralized with sodium counter ions. Additionally, a 600 mM KCl salt buffer was added to the system, replicating the experimental conditions. First, we conducted a 1 ns NVT equilibration step, followed by a 1 ns NPT equilibration step, while applying position constraints to the C-α atoms. Thereafter, the system was simulated for 20 ns in an NPT ensemble in order to equilibrate the protein structure.

The SMD simulations were carried out using the final structure obtained from the equilibration simulation. The C-α atom positions of cysteine residues 319 and 521 served as the collective variables in our SMD simulation. This is achieved by applying an umbrella potential to the C-α atoms with a spring constant of 1100 pN/nm, mimicking the effect of the DNA handles on the protein^48^. One of the C-α atoms is fixed in position, while the other is pulled along the x-axis at a constant velocity. During the simulation, the forces on the C-α atoms and their positions were saved every 200 ps.

During the SMD simulations, a pulling velocity of 1·10^−4^ nm/ps (i.e., 0.1 m/s) was used. The chosen velocity strikes a balance between approximating the experimental pulling velocity and maintaining a reasonable simulation cost. However, it is important to note that the experimental pulling velocities are still outside of the scope of MD simulations for our application, as they are still six orders of magnitude slower.

Generally, SMD simulations can be quite computationally expensive due to the need for a large simulation box^49^. The box size is necessary to fully solvate the protein throughout the entire extension, which results in a high number of solvent molecules. In order to improve the efficiency of the SMD simulation and therefore decrease the attainable pulling velocity, we employed a constant cross-section telescoping box^50^. The telescoping box scheme refers to splitting up the SMD simulation into stages, where in between the stages, the simulation box is extended along the pulling axis. The purpose of this scheme is to enhance the numerical efficiency by decreasing the number of solvent molecules.

As the first step in the SMD simulation, we placed the equilibrated protein structure in a triclinic simulation box. Here we aligned the vector that connects the cysteine residues 319 and 521 along the pulling axis and ensured that there is a 1 nm distance between the periodic images of the protein. As previously done, we neutralized the system with sodium counter ions after solvating the protein and subsequently added a 600 mM KCl salt buffer. We prepared an equilibrated water box that has the same yz-cross section and a length of 10 nm along the x-axis, which was concatenated to our system along the pulling axis. After merging the solvated protein with the equilibrated water box, position restraints are applied to the protein, and the solvent is allowed to be equilibrated for 100 ps under an NVT simulation. After the brief solvent equilibration, the pulling simulation is started for 100 ns under NVT conditions, which extends the Cys-Cys distance by 10 nm along the pulling axis.

The procedure of extending the simulation box and pulling the protein into the newly available solvent is repeated until all the secondary and tertiary structure motifs of the protein are fully unfolded. Based on the predicted contour length change of 71.8 nm by completely unfolding OpuAC from a natively folded state, we needed to perform this procedure at most 8 times. At the start of each unfolding simulation, approximately 35,000 atoms were present in the system, while the final system contained more than 430,000 atoms. Practically this means that the application of telescoping simulation boxes reduced our computational cost by 40%. The SMD simulation was replicated seven times for statistical validation, resulting in more than 4.5 μs of simulation time.

#### Simulation Data Analysis

During SMD simulations, it was observed that the OpuAC protein follows a distinct folding pathway. The first step involves the unraveling of the N-terminal beta sheet from lobe 1. Following this, the rest of lobe 1’s structure unfolds, and eventually, lobe 2 also unravels (**Figure 3B**).

Although the order of this unfolding pathway is conserved over all replicas, variations on the unfolding pathway have been identified. During the initial two replica simulations, it was observed that the N-terminus exhibited strong interaction with the unfolded residues of lobe 1 through the creation of transient beta sheets. These short-lived beta sheets caused a significant increase in the amount of force required to fully unfold lobe 1, reaching approximately 700 pN. Additionally, replica 7 exhibits an anomalous unfolding pathway where a section of lobe 1, which is still folded, becomes separated from the protein as a whole. This domain is made up of two beta sheets and an alpha helix, which requires a considerable amount of force to completely unfold during the simulation’s later stages. In comparison to the other replicas, this stable subdomain causes the protein to only unfold completely after a distance of 63 nm is reached, while in the other replicas, all significant structures unfolded before 60 nm.

During the SMD simulations, the measured forces on the C-α atoms of cysteine residues 319 and 521 were recorded every 10000 steps (i.e., 20 ps). Due to thermal fluctuations in the SMD simulations, there is a significant background noise on these measured forces. To reduce this background noise and isolate the contribution of the protein unfolding, we applied a Savitzky-Golay filter with a window length of 1000 measurements and a degree 3 polynomial^51^.

The seven replicate FD curves resulting from the MD simulations are displayed in the left-hand side panel of **Figure S6**. The individual simulation windows of the telescoping box scheme were overlaid on the measurements using the shaded areas. At each transition between windows, the simulation is halted, and the periodic box is extended in the pulling direction by adding solvent. When comparing the simulated and experimental FD curves, it was observed that there was a significant difference in the measured forces by an order of magnitude. This discrepancy can be attributed to the much faster pulling velocity employed in the SMD simulations.

The average FD curve is shown in the right-hand side panel of **Figure S6**. The overlaid vertical lines represent the unfolding of the mechanical intermediates identified in the MD simulations. The measured contour length changes between the intermediates show a qualitative agreement between the experiments and simulations. One notable distinction between the experimental and simulated FD curves is the absence of the unfolding of the fourth intermediate (I4) in the simulated FD curves. This we attribute to the much faster pulling speed and the associated high forces in the simulation, combined with the relatively low mechanostability of I4. It should be noted that the standard error on the average FD curve during the I1→I2 unfolding transition is relatively high, which can be attributed to the formation of transient beta sheets by the N-terminus. As previously mentioned, the formation of beta sheets between unfolded residues and the N-terminus increases the force required to unfold lobe 1.

