## Supplemental Data and Figures for "Probing the stability and interdomain interactions in the ABC transporter OpuA, using single-molecule optical tweezers"

### Supplementary Tables and Figures

| 50 mM HEPES-K pH 7.0, 600 mM KCl | | | | | |
|---|---|---|---|---|---|
| ***Transition*** | **Counts** | **AverageL$_c$ (nm)** | **SD** | **Average unfolding force (pN)** | **SD** |
| *Y-X* | 5 | 15.0 | 6.34 | 5.8 | 2.03 |
| *X-A* | 23 | 13.9 | 1.40 | 16.1 | 4.46 |
| *A-B* | 11 | 14.3 | 3.89 | 19.2 | 5.88 |
| *B-C* | 10 | 25.5 | 1.79 | 18.1 | 5.20 |
| *C-D* | 11 | 19.0 | 2.92 | 18.3 | 5.46 |
| *D-E* | 6 | 14.8 | 2.09 | 20.8 | 4.17 |
| *A-C* | 7 | 41.3 | 6.00 | 17.7 | 6.19 |
| *B-D* | 2 | 43.4 | 2.50 | 21.1 | 2.60 |
| *X-B* | 2 | 27.6 | 1.15 | 17.3 | 1.35 |

| 50 mM HEPES-K pH 7.0 | | | | | |
|---|---|---|---|---|---|
| ***Transition*** | **Counts** | **Average ΔL$_c$ (nm)** | **SD** | **Average unfolding force (pN)** | **SD** |
| *Y-X* | 9 | 11.0 | 5.38 | 12.8 | 5.85 |
| *X-A* | 27 | 11.3 | 1.67 | 21.8 | 4.44 |
| *A-B* | 12 | 14.2 | 2.27 | 20.6 | 5.36 |
| *B-C* | 9 | 25.5 | 3.41 | 18.4 | 2.81 |
| *C-D* | 16 | 16.6 | 2.70 | 20.4 | 3.64 |
| *D-E* | 9 | 14.8 | 1.97 | 26.2 | 4.54 |
| *A-C* | 6 | 37.7 | 2.08 | 18.4 | 2.87 |
| *A-D* | 4 | 56.2 | 4.61 | 28.0 | 4.66 |
| *B-D* | 5 | 41.2 | 2.85 | 23.3 | 2.52 |
| *X-B* | 4 | 24.9 | 0.81 | 19.2 | 3.24 |
| *X-C* | 1 | 53.1 | - | 30.0 | - |

***Table S1: Contour length changes and unfolding forces of OpuA.*** *Average and standard deviation of the change in contour length (ΔL$_c$) and unfolding force for every transition, related to Figure 5. Additionally, average lengths obtained from transitions two intermediates unfolded simultaneously within the time resolution of the experiment (e.g., from A to C) are shown but they have not been used for the statistics in **Figure 5**. Data has been obtained from 4 molecules, 34 unfolding traces (50 mM HEPES-K pH 7.0, no extra salt) and 6 molecules, 28 unfolding traces (50 mM HEPES-K pH 7.0, 600 mM KCl).*

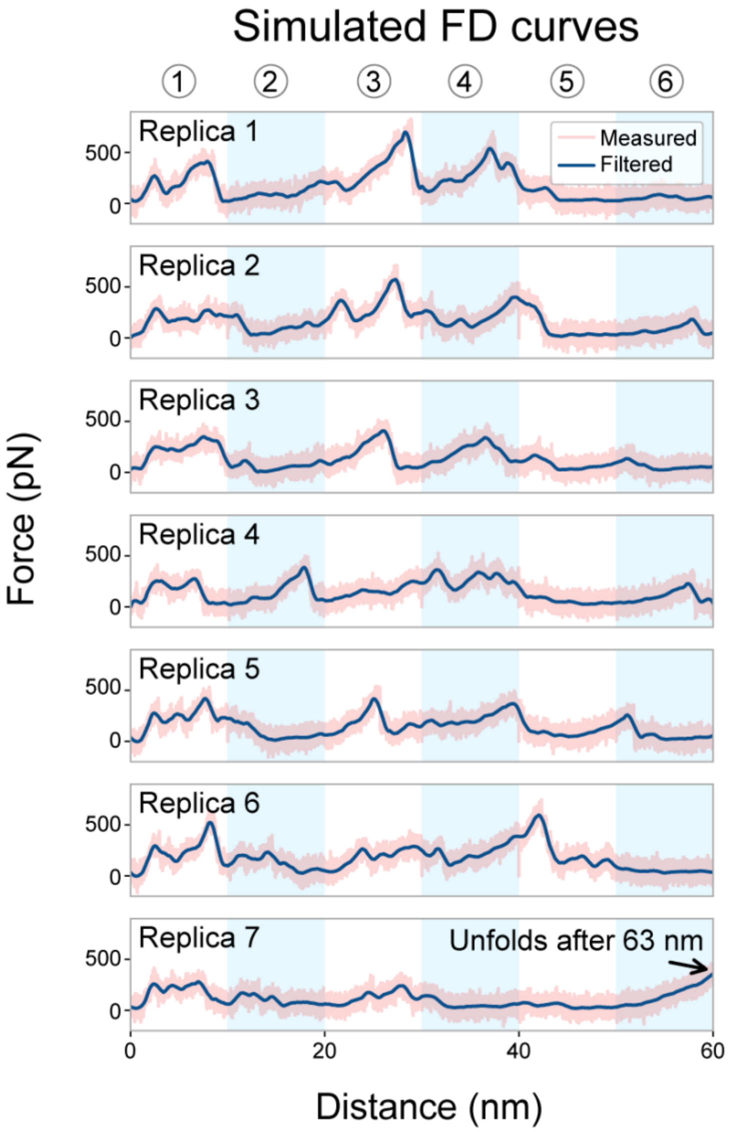

***Figure S1: Force distance curves measured during the MD simulations, related to Figure 3.*** *The unprocessed data obtained from the seven replica simulations and their corresponding filtered traces are shown. In this panel, the vertically shaded areas indicate the windows of the telescoping box scheme.*

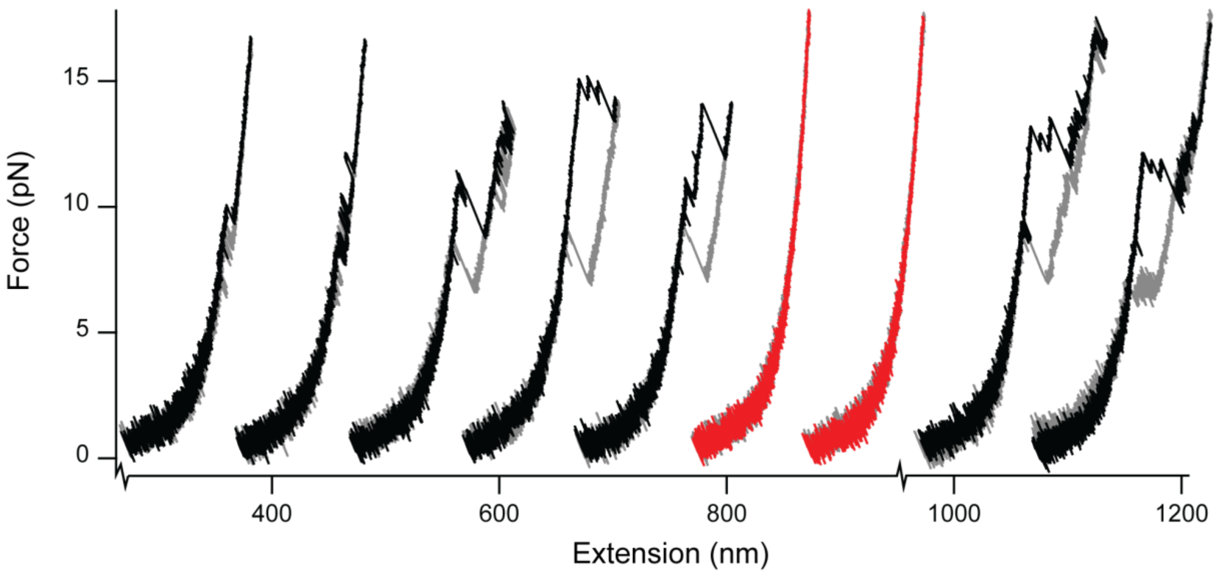

***Figure S2: Force distance curves of OpuA in 50 mM HEPES-K pH 7.0, 600 mM KC, related to figure 5.*** *A sequence of force-distance where both unfolding (black) and misfolds without unfolding (red) have been observed at 20 nm/s. Folding curves are shown in gray with a horizontal offset of 100 nm.*

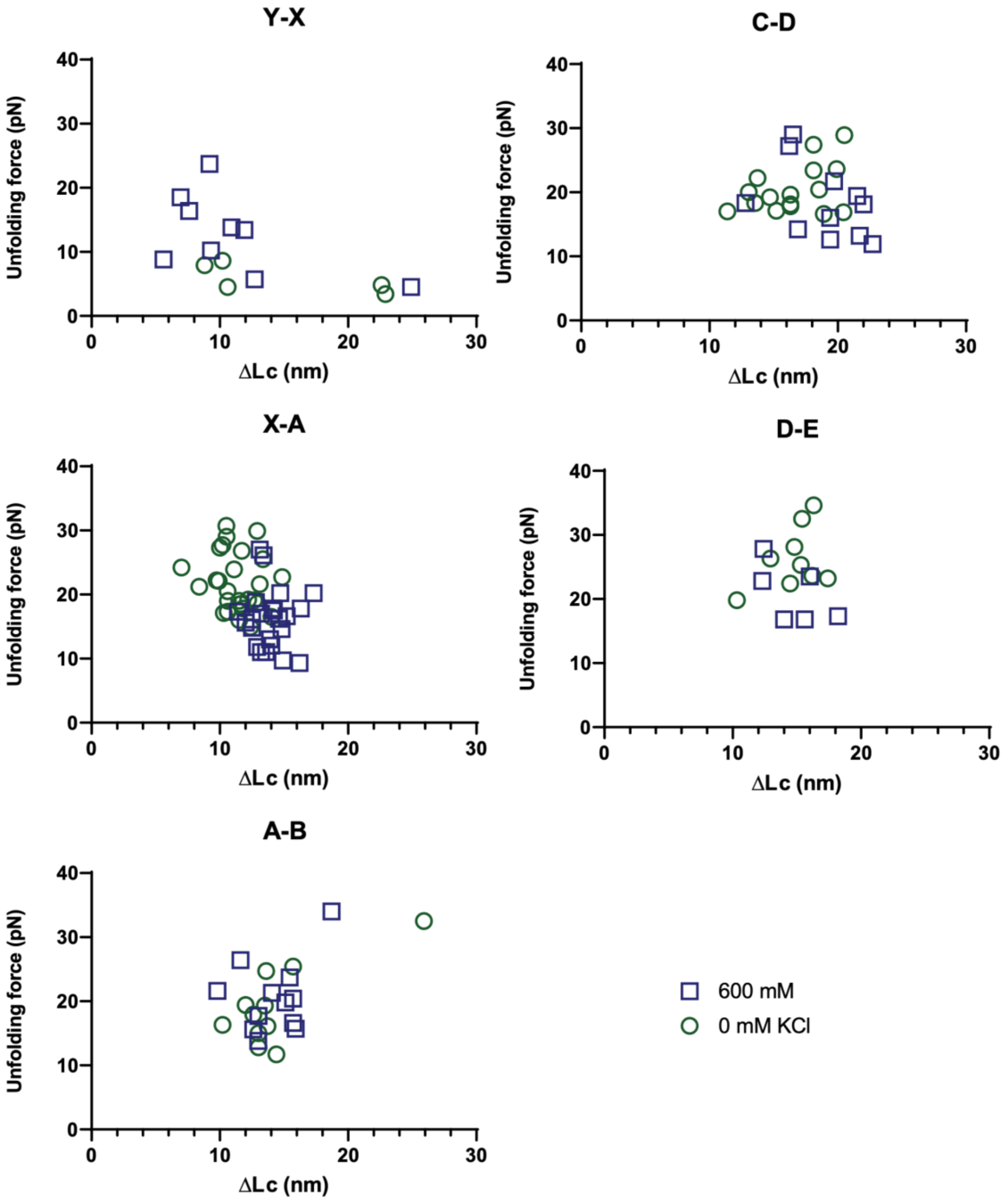

***Figure S3: Scatterplots of the unfolding forces and gains in contour length ($\Delta L_c$) of the transitions observed in OpuA, related to Figure 5.*** *Data points have been collected from 4 molecules, 34 traces (50 mM HEPES-K pH 7.0, no extra salt) and 6 molecules, 28 traces (50 mM HEPES-K pH 7.0, 600 mM KCl). Traces have been collected at 20 nm/s with a waiting time of 1 s.*

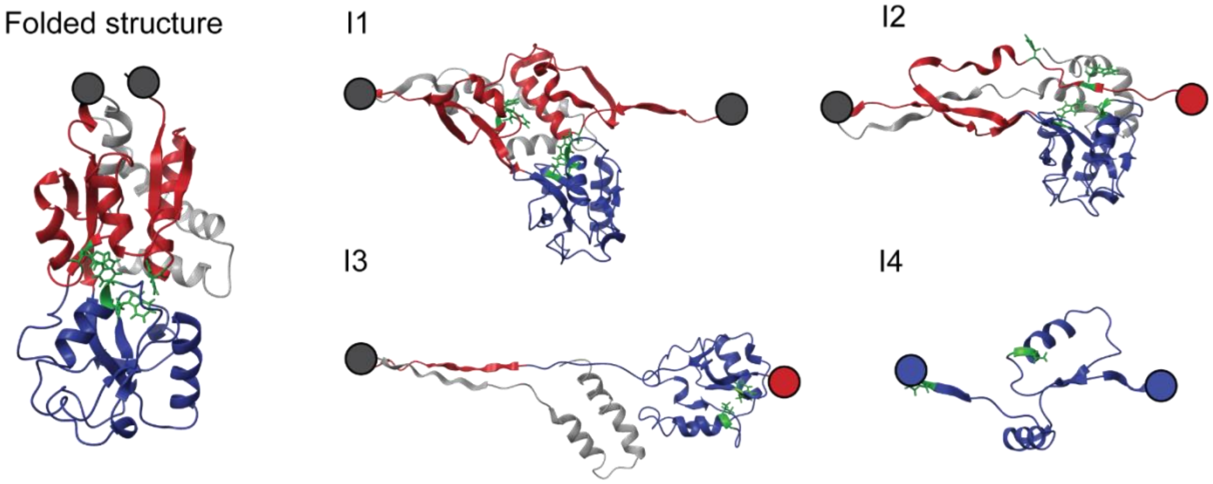

***Figure S4. Structures of the mechanical intermediate states found by MD simulations based on the crystal structure, related to Figure 3.*** *Colors in the structure (PDB: 3L6G) represent lobe 1 (red), lobe 2 (blue) and the C-terminal tail (light gray) which is not probed by the optical tweezers experiment. The drawn circles represent the attachment point of the handle (dark gray) and a fully unfolded amino acid chain corresponding to (part of) lobe 1 (red) and/or lobe 2 (blue), which has been cropped for simplicity of the figure. Residues involved in glycine betaine binding have been highlighted in green.*
